# Dynamic integration and segregation of amygdala subregional functional circuits linking to physiological arousal

**DOI:** 10.1101/2020.11.21.392910

**Authors:** Yimeng Zeng, Fuxiang Tao, Zaixu Cui, Liyun Wu, Jiahua Xu, Wenshan Dong, Chao Liu, Zhi Yang, Shaozheng Qin

**Affiliations:** State Key Laboratory of Cognitive Neuroscience and Learning, Beijing Normal University, Beijing, China; IDG/McGovern Institute for Brain Research, Beijing Normal University, Beijing, China; Beijing Key Laboratory of Brain Imaging and Connectomics, Beijing Normal University, Beijing, China; Chinese Institute for Brain Research, Beijing, China; School of Psychology, South China Normal University, Guangzhou, China; Shanghai Key Laboratory of Psychotic Disorders, Shanghai Mental Health Center, Shanghai Jiao Tong University School of Medicine, Shanghai, China

**Keywords:** Amygdala, fMRI, dynamic states, arousal, emotion

## Abstract

The dynamical organization of brain networks is essential to support human cognition and emotion for rapid adaption to ever-changing environment. As the core nodes of emotion-related brain circuitry, the basolateral amygdala (BLA) and centromedial amygdala (CMA) as two major amygdalar nuclei, are recognized to play distinct roles in affective functions and internal states, via their unique connections with cortical and subcortical structures in rodents. However, little is known how the dynamical organization of emotion-related brain circuitry reflects internal autonomic responses in humans. Using resting-state functional magnetic resonance imaging (fMRI) with K-means clustering approach in a total of 79 young healthy individuals (cohort 1: 42; cohort 2: 37), we identified two distinct states of BLA- and CMA-based intrinsic connectivity patterns, with one state (integration) showing generally stronger BLA- and CMA-based intrinsic connectivity with multiple brain networks, while the other (segregation) exhibiting weaker yet dissociable connectivity patterns. In an independent cohort 2 of fMRI data with concurrent recording of skin conductance, we replicated two similar dynamic states and further found higher skin conductance level in the integration than segregation state. Moreover, machine learning-based Elastic-net regression analyses revealed that time-varying BLA and CMA intrinsic connectivity with distinct network configurations yield higher predictive values for spontaneous fluctuations of skin conductance level in the integration than segregation state. Our findings highlight dynamic functional organization of emotion-related amygdala nuclei circuits and networks and its links to spontaneous autonomic arousal in humans.

## 1. Introduction

The human brain, as an open and nuanced system, spontaneously abounds with its dynamic nature. Such spontaneity is thought to meet ever-changing environmental needs vital to survive, involving in a massive amount of information flows between the brain and body. Spontaneous activity of functional brain networks does not remain constant, but undergoes dynamic fluctuations over time (Bressler et al., 2010; Bassett et al., 2011; Braun et al., 2015) that are thought to reflect affective and/or homeostatic states along with autonomic and hormonal responses (Dolan, 2002; Deco et al., 2011; Calhoun et al., 2014). Dysregulated intrinsic dynamics of brain networks critical for emotion and related behaviors have been linked to mood and anxiety disorders as well as among other psychiatric conditions (Sripada et al., 2012; Bickart et al., 2014; Rashid et al., 2016). The amygdala lies at the core of emotion-related brain networks to broadcast affective state information across widespread brain regions in (para)limbic systems, emotional and salience networks critical for autonomic arousal, salience detection, emotion perception and regulation (Davis & Whalen, 2001; LeDoux, 2003; Seeley et al., 2007; Pessoa & Adolphs, 2010). However, little is known about the dynamic nature of the amygdala and its functional circuits, and even less is known whether and how their dynamic features are associated with internal autonomic states in humans.

The amygdalar complex encompasses multiple anatomical subregions with distinct connections that support distinct affective functions (LeDoux, 2000; LeDoux, 2007). The basolateral amygdala (BLA) and centromedial amygdala (CMA) are two major groups of amygdalar nuclei that form dedicated networks for distinct functions via their unique pattern of interactions with other cortical and subcortical structures (LeDoux, 2000; LeDoux, 2007). These structures include the insular complex critical for autonomic arousal, visceral sensations, and interoceptive processing (Craig, A. D., 2009; Pessoa & Adolphs, 2010), the anterior cingulate cortex responsible for salience detection and executive attention (Critchley, H. D., 2005), the inferior temporal cortices responsible for perceptual processing of emotional stimuli (Hermans et al., 2014), and the lateral and orbital prefrontal cortex critical for regulation of emotions (Machado et al.,2009). Within the past decade, there has been great progress in functional neuroimaging techniques that allows researchers to map spontaneous functional coupling of the amygdalar nuclei with large-scale brain networks in humans (Roy et al., 2009; Salzman et al., 2010; Qin et al., 2012; Sladky et al., 2015). Seed-based correlation analysis of spontaneous brain activity has been the mainstay of conventional approaches for examining intrinsic functional organization of large-scale brain networks including amygdala-based emotional networks (Roy et al., 2009; Uddin et al., 2009). This provides useful information to identify specific target networks with the BLA and CMA, but offers insufficient insight into how these networks dynamically organize over time to regulate the brain’s affective functions. Thus, novel approaches from a dynamic and nonstationary systems perspective are required to mitigate this limitation. Beyond simple scan-length averages (so-called static methods), recent emerging innovative approaches by capturing time-varying properties of connectivity from a systems level have begun to illustrate the dynamic nature of spontaneous brain activity across large-scale networks (i.e., brain state) in humans (Hutchison et al., 2013; Calhoun et al., 2014). However, the dynamical organization of functional brain networks associated with the amygdalar nuclei still remains open.

There is now increasing evidence converging onto that the dynamic nature of brain functional organization is regulated by the ascending neuromodulatory systems such as locus coeruleus (LC) releasing noradrenaline by acting on widespread brain networks to alter the neuronal excitability and drive the integration of distributed neurons (Aston-Jones & Cohen, 2005). Indeed, the dynamics of network topology has been linked to autonomic arousal levels varying as a function of firing patterns of LC neurons (Aston-Jones & Cohen, 2005; Eldar et al., 2013; Shine et al., 2018). In particular, the amygdalar nuclei, including BLA and CMA, constitute a large amount of neurotransmitter receptors which could transiently modulate neuronal excitability of the information processing through their projections to specific target circuits and networks (LeDoux, 2007). Critically, one recent study in rodents shows that amygdala neuronal ensembles dynamically encode affective or homeostatic internal states, through two major populations of neurons of the basal nucleus of the amygdala that widely broadcasts internal state information via several output pathways to larger brain networks (Gründemann et al., 2019). Despite differences in neuroanatomical and neurochemical properties of the amgydala nuclei between rodents and humans (Pabba et al., 2013), recent studies from human functional neuroimaging have also demonstrated a link between the brain’s intrinsic functional activity and physiological fluctuations of autonomic arousal. For instance, connectivity changes in the amygdala and dorsal anterior cingulate cortex (dACC) with a set of regions including brainstem, thalamus and putamen as well as dorsolateral prefrontal cortex appear to covary with heart rate variability (HRV) (Chang et al., 2013). Spontaneous activity in the posterior cingulate cortex (PCC) of default mode network (DMN) and the anterior cingulate cortex (ACC) and anterior insular (AI) of task-positive network (TPN) also covaries with non-specific skin conductance response at resting state (Fan et al., 2012). Recently, a study observed dynamic fluctuations of amygdala functional connectivity relevant to physiological arousal observed in a fear conditioning paradigm (Baczkowski et al., 2017). Yet, how the dynamical organization of large-scale networks associated with the amygdalar nuclei reflects internal autonomic states in humans remains unknown.

Here we addressed the above open questions by investigating time-varying connectivity patterns of functional circuits associated with the two major amygdalar nuclei (i.e., BLA, CMA) and their links to internal autonomic responses in a total of 79 young healthy adults across two independent cohorts. Using resting-state (8-min) functional magnetic resonance imaging (fMRI), we first examined time-varying intrinsic functional connectivity properties of the BLA and CMA and their network configurations in Cohort 1 (N = 42). The prior observer-independent cytoarchitectonically determined probabilistic maps were used to define the BLA and CMA masks (Amunts et al., 2005; Eickhoff SB, et al. 2005). The conventional sliding-window approach was implemented to identify time-varying spontaneous functional connectivity of coupling and connected networks of regions (Sakoğlu et al., 2010; Allen et al., 2014; Chen et al., 2016). K-means clustering method, as one of the unsupervised machine-learning algorithms, was used to identify dissociable connectivity states that are referred as time-varying BLA- and CMA-based intrinsic functional network configurations among multiple brain regions. Resting-state fMRI (8-min) data from a second independent cohort of 37 participants with higher spatial resolution (see Methods) were used to gain a better localization of the amygdala nuclei and ensure the reproducibility and robustness of the observed dynamic states. We further implemented Elastic-net regression to examine whether time-varying connectivity patterns of the amygdalar nuclei are predictive of automimic arousal measured by skin conductance levels. Based on the neurophysiological models of network dynamics and autonomic arousal (Critchley, H. D., 2005; Young et al., 2017; Shine et al., 2018), we predict that intrinsic functional connectivity patterns associated with the two major amygdala nuclei would undergo dissociable time-varying states and such dynamics would predict spontaneous autonomic responses.

## 2. Methods and materials

### 1.1. Participants

This study included a total of 79 young healthy participants from two independent cohorts. The first cohort consisted of 42 young healthy participants (mean age ± SD: 22.62 ± 0.99 years ranged from 21 to 24, 27 females) after dropping out invalid participants because of excessive head motion (**Cohort 1**). A second independent cohort of 37 adults matched in age and sex (mean age ± SD: 22. 08 ± 1.65 years ranged from 20 to 25, 18 female) was recruited to undergo fMRI with the concurrent recording of skin conductance (**Cohort 2**). All of the participants reported no history of any neurological or psychiatric disorders, and no current use of any medication or recreational drugs. The experiment and procedures were approved by the Institutional Review Board for Human Subjects at Beijing Normal University in accordance with the standards of the Declaration of Helsinki. Written, informed consent was obtained from all participants before the experiment.

### 1.2. Imaging acquisition

For **Cohort 1**, brain images were acquired from a Siemens 3T scanner (Siemens Magnetom Prisma syngo MR D13D, Erlangen, Germany) with a 64-channel phased-array head coil in Peking University. Participants were instructed to keep their eyes open and remain still for a period of 8-min resting-state fMRI scan. T2-weighted images were recorded using an echo-planar imaging (EPI) sequence, with 177 volumes (axial slices, 33; slice thickness, 3.5 mm; volume repetition time, TR, 2000 ms; echo time, TE, 30 ms; flip angle, 90; voxel size, 3.5 × 3.5 × 3.5 mm; field of view, FOV, 224 × 224 mm). High-resolution anatomical T1-weighted images were collected using three-dimensional sagittal T1-weighted magnetization-prepared rapid gradient echo (MPRAGE) sequence (192 slices; TR, 2530 ms; TE, 2.98 ms; slice thickness, 1 mm; flip angle, 7°; voxel size, 0.5 × 0.5 × 0.5 mm; FOV, 256 × 256 mm).

For **Cohort 2**, whole-brain imaging data were collected on a 3T Siemens Prisma MRI system in Peking University. Participants were instructed to keep their eyes open and remain still for a period of 8-min resting-state fMRI scan. Functional images were collected using a multi-band echo-planar imaging (mb-EPI) sequence (slices, 64; slice thickness, 2 mm; TR, 2000 ms; TE, 30 ms; flip angle, 90°; multiband accelerate factor, 2; voxel size, 2.0 × 2.0 × 2.0 mm; FOV, 224 × 224 mm). Structural images were acquired through three-dimensional sagittal T1-weighted magnetization-prepared rapid gradient echo (MPRAGE) sequence (192 slices; TR, 2530 ms; TE, 2.98 ms; slice thickness, 1 mm; voxel size, 1.0 × 1.0 × 1.0 mm, interpolated to 0.5 × 0.5 × 0.5 mm; flip angle, 7°; inversion time, 1100 ms; FOV, 256 × 256 mm). Field map images were acquired with grey field mapping sequence (64 slices; TR, 635 ms; TE1, 4.92 ms; TE2, 7.38 ms; slice thickness, 2 mm; voxel size, 2.0 × 2.0 × 2.0 mm; flip angle, 60°; FOV, 224 × 224 mm).

### 1.3. Imaging preprocessing

Functional imaging preprocessing was performed using tools from Statistical Parametric Mapping SPM12 (http://www.fil.ion.ucl.ac.uk/spm). For **Cohort 1**, the first five volumes were discarded for signal equilibrium. Images were then firstly corrected for distortions related to magnetic field inhomogeneity. Subsequently, these functional images were realigned for rigid-body motion correction and corrected for slice acquisition timing. Each participant’s images were then co-registered to each participant’s gray matter image segmented from corresponding high-resolution T1-weighted image, spatially normalized into a standard stereotactic Montreal Neurological Institute (MNI) space and resampled into 2-mm isotropic voxels. Finally, images were smoothed by an isotropic three-dimensional Gaussian kernel with 6 mm full-width at half-maximum.

### 1.4. Skin conductance (SC) recording and analysis

Skin conductance data in **Cohort 2** (N = 37) were recorded simultaneously with fMRI scanning using an MRI-compatible Biopac MP 150 System (Biopac, Inc., Goleta, CA). Two Ag/AgCl electrodes filled with isotonic electrolyte medium were attached to the center phalanges of the index and middle fingers of the left hand, connecting to a Biopac GSR100C module. The gain set to 5, and the high pass filters set to DC. Data were acquired at 1000 samples per second.

### 1.5. Skin conductance level (SCL) preprocessing

SCL data from all participants were first inspected to ensure complete SCL recording and exclude incomplete SCL recordings due to malfunctioning of machine or missing of start(end) triggers when scanning. Second, we excluded participants with relatively large motion artifacts and other related factors that could distorted the quality of SCL data. Next, according to the procedures by previous studies (Boucsein et al., 2012; Braithwaite et al., 2013), we conducted five sequential operations as follows: 1) A cut off low pass filter at 10 Hz to remove high frequency noise, 2) A detrending operation to move the slow-drift, 3) Down-sampling the SCL by averaging data points within each 2 sec, 4) Each data normalized by dividing its standard deviation, 5) Window-averaged SCL time points in corresponding to each state-wise functional connectivity. Finally, to avoid potential subjectivity when determining exclusion SCL time series, we also conducted additional analyses by including 7 participants who were excluded according to the second procedure mentioned above. According to our research questions and above processing procedures, our present study focused on a relatively slow, tonic-like component of skin conductance level (SCL) fluctuations rather than phasic component of skin conductance response (SCR).

### 1.6. Regions of interest (ROIs) definition

Four ROIs encompassing the BLA and CMA for each hemisphere were created separately using cytoarchitectonically defined probabilistic maps of the amygdala. Maximum probability maps were used to create nonoverlapping amygdala subregions using the Anatomy Toolbox (Eickhoff SB, et al. 2005). Voxels were included in the maximum probability maps only if the probability of their assignment to the BLA or CMA was higher than any other nearby structures. Each voxel was exclusively assigned to only one region. Overlapping voxels were assigned to the region that had the greatest probability, resulting in two nonoverlapping ROIs representing CMA and BLA subregions for each hemisphere. Based on previous studies (Roy et al., 2003; Qin et al., 2012), we selected the five target networks of interest defined in the AAL atlas (**Fig. 1D**). These target networks include (1) subcortical structures including striatum, thalamus and midbrain; (2) crus and vermis/declive regions of the cerebellum; (3) uni- and polymodal association cortex including sensory and motor areas that consist of visual cortex (VC), parahippocampal gyrus (PHG), inferior temporal cortex (ITC), middle (MTG) and superior temporal gyrus (STG), premotor cortex (PMC) and sensorimotor cortex (SM); (4) limbic and paralimbic structures including hippocampus (Hipp), insula (Ins), middle (mCC) and posterior (PCC) portions of cingulate cortex; and (5) prefrontal cortex including anterior cingulate cortex (ACC), middle (MFG) and inferior frontal gyrus (IFG), and medial prefrontal cortex (MPFC). The rationale of selection for these networks of interest are depicted in **Table. S1**.

**Fig. 1.**
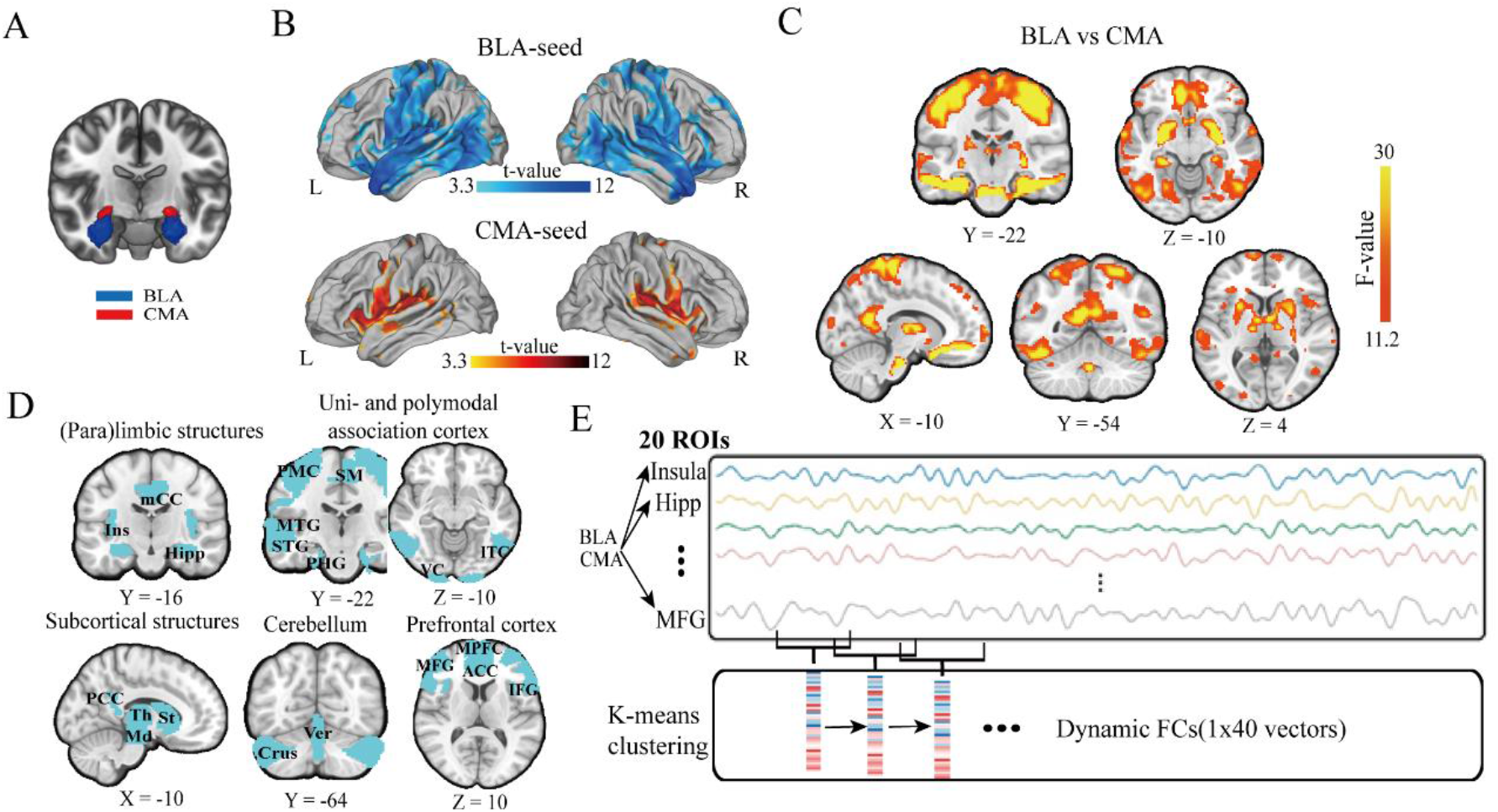
An overview of time-varying intrinsic functional connectivity of BLA- and CMA-based target networks by sliding window and K-means clustering. (**A**) A coronal view of the BLA (blue) and CMA (red) seeds. (**B**) Lateral views of significant clusters in widespread brain regions showing intrinsic functional connectivity with the BLA and CMA seeds. (**C**) Brain regions showing significant clusters exhibiting the main effects of BLA vs. CMA. (**D**) Representative views of the anatomically defined five target networks of interest. (**E**) An illustration of K-means cluster analysis. Connectivity between BLA/CMA seed regions and 20 target regions from five networks of interest were computed in each window. Then, K-means clustering was conducted on window sequence. Notes: L, left; R, right; BLA, basolateral amygdala; CMA, centromedial amygdala; mCC, middle portions of cingulate cortex; PCC, posterior portions of cingulate cortex; VC, visual cortex; ITC, inferior temporal cortex; MTG, middle temporal gyrus; STG, superior temporal gyrus; PMC, premotor cortex; SM, sensorimotor cortex; ACC, anterior cingulate cortex; IFG, inferior frontal gyrus; MPFC, medial prefrontal cortex; MFG, middle frontal gyrus.

### 1.7. Functional connectivity analysis

Time series of each seed (averaging across all voxels within ROI) were filtered with a bandpass temporal filter (0.008 to 0.10 Hz) and extracted. Six motion parameters, cerebrospinal fluid and white matter of each participant that account for potential physiological noise and movement-related artifacts were regarded as covariates of no interest. Subsequently, the resultant time series were demeaned. For each participant, two separate functional connectivity analyses were performed for both BLA and CMA seed separately and significant clusters were determined using a height threshold of *p* < 0.001 and an extent threshold of *p* < 0.05 corrected for multiple comparisons. To further investigate regions exhibiting different functional connectivity with BLA and CMA-seed, the contrast parameter images for each of the four seed ROIs from the individual level analyses were submitted to a second-level group analysis by treating participants as a random variable in a 2-by-2 ANOVA with amygdala subregions (BLA vs. CMA) and hemispheres (left vs. right) as within-subject factors. Significant clusters were determined using a height threshold of *p* < 0.001 and an extent threshold of *p* < 0.05 corrected for multiple comparisons. For later K-means analyses, target ROIs of each seed were defined by the combination of significant clusters exhibiting main effects of BLA vs CMA in ANOVA analyses and 20 regions of interest derived from anatomically-defined AAL atlas. Subsequently, both times series of each seed and associated target ROIs were extracted by the same procedure mentioned above. And Pearson’s correlation coefficients, representing the strength of functional connectivity of each seed with corresponding target masks, were computed between amygdala subregion-seeded time series and target time series.

For replication purpose, we also defined 20 ROIs using an independent finer-grained AICHA atlas (Joliot et al., 2015), except for three ROIs locating at the midbrain, crus and vermis. Because there are no corresponding parcellations for these three ROIs in the AICHA atlas and we simply used these three ROIs defined by the AAL atlas. Notably, all target ROIs were implemented without combination in replication analyses to test robustness of state clustering results. Thereafter, parallel analyses were conducted for our fMRI data from **Cohort 1** and **Cohort 2**.

### 1.8. Time-varying functional connectivity analysis and K-means clustering

Time-varying functional connectivity was assessed by using a widely used sliding-window approach (Sakoğlu et al., 2010; Bassett et al., 2011; Allen 2014; Shine et al., 2016) in MATLAB R2015b (MathWorks, Natick, USA). The entire temporal time series were divided into a series of windows with a specific window length (α = 40 TR) and step (β = 1 TR). The total quantity of windows was defined by a parameter *W*,

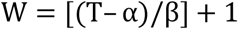

where *T* is a total number of time points. Initial point *S* for each window *i* was defined as:

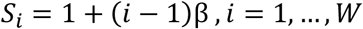

while endpoint *E* was defined as:

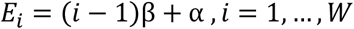

For state clustering, a brain state here is referred to specific spatial configurations of time-varying intrinsic functional connectivity of the BLA and CMA seeds with 20 target ROIs defined above. We first computed Pearson’s correlation coefficients between BLA and CMA seed and associated target ROIs within each time window. Each participant’s BLA- and CMA-seed functional connectivity within each time window was then transformed into one-dimensional vectors. Next, K-means clustering of windowed correlation matrices was implemented to identify different states of dynamics of BLA- and CMA-based intrinsic functional connectivity. The algorithm was repeated with 1000 random iterations to exclude the sensitivity of K-means to initial conditions.

A paramount step of clustering analysis is determining the optimal parameter set (K, α, β). Various parameter sets were attempted to fulfill this goal (*K* = 2 − 15, sliding window length α = 30 − 50 TRs, step β = 1.0 − 3.0 TRs). The optimal parameter sets were obtained after evaluation of silhouette coefficient (Rousseeuw, 1987) and CalinskiHarabasz criterion (Caliński & Harabasz, 1974), in which higher value indicates larger cohesion and less dispersity.

### 1.9. Time-lagged cross-correlation (TLCC) analysis for dynamic states

We implemented time-lagged cross-correlation (TLCC) to investigate temporal coherence between BLA and CMA intrinsic target networks in different states (Adhikari et al., 2010). We first concatenated all of time-varying windows for BLA- and CMA-based intrinsic connectivity with 20 target ROIs in each state across participants. We then averaged window-wise series of BLA- and CMA-based intrinsic connectivity separately, for each state. Thereafter, we calculated TLCC between averaged BLA and CMA connectivity windows with a lag range from −40 to 40. To calculate TLCC on finer-grained level, we further divide whole window series of averaged BLA and CMA-based connectivity into epochs, with each epoch length setting at 80. Then, TLCC was computed under each epoch separately with the same lag range from −40 to 40. We define the result based on this procedure as TLCC (finer-grained).

### 1.10. Machine learning-based prediction analysis

To examine whether dynamic states, featured by their time-varying functional networks of BLA- and CMA seeds, can be robustly linked to ongoing SCL fluctuations, two Elastic-net regression models with 10-fold cross validation were implemented for detected states separately. For each model, a total of 40 dynamic connections were extracted as a feature vector (i.e., 1× 40 vector) and corresponding window-average SCL extracted as a response variable at each temporal window. The Statistics and Machine Learning Toolbox in MATLAB 2015a (MathWorks, Natick, USA) was used to perform the Elastic net regression (Boyd S., 2010). Elastic-net model earns the benefits of excluding little-contributed variables while reducing overfitting problem by implementing L1 and L2-regularization in the model (controlled by parameter *α* and *λ*). To determine the optimal parameter set (*α, λ*) for the Elastic-net regression model, we applied a grid search with *α* chosen from 20 values in the range of [0.05, 1.0] and *λ* chosen from 100 values using a default lambda sequence pre-defined in Statistics and Machine Learning Toolbox. Then, 10-fold cross-validation was applied to examine the performance of Elastic-net regression at different parameter settings. Finally, (*α, λ*) with minimum cross-validation error plus one standard deviation was select for later analysis (Tsankov et al., 2015).

To determine the significance of predictive accuracy for Elastic-Net regression models, we employed a stringent permutation test to account for temporal autocorrelations present in spontaneous BOLD signal fluctuations and SCL time series (Kauppi et al., 2010). For each state, we circularly shifted both window-wise BLA/CMA-based connectivity series within each state and corresponding SCL time series in the same state in a random manner, on an individual level. We then trained Elastic-net regression models using the permuted data across participants on a group level. The total number of permutation times for each state was set to 5000. The *p* values of the mean correlation *r* were calculated by dividing the number of permutations that showed a higher value than the actual value for the real sample by the total number of permutations. To test the difference of prediction accuracy for SCL among states, a bootstrapping method was implemented. Specifically, we conducted repeated resampling from time-varying connectivity vectors and their corresponding window-wise SCL value for each state separately. Then, we trained Elastic-net models under bootstrapped data with the same hyperparameters settings in **Fig. S10**. Next, Pearson’ correlation was implemented to estimate the correlation between predicted SCLs and observed SCLs. We thus created the simulated distribution of Pearson’s correlation between observed and predicted SCLs for each state respectively. Finally, two sample t tests were employed to compare the differences of distributions (Fisher-z transformed). The resample times for bootstrapping were set to 5000 and the population size for each resample was set to 1500. Finally, prediction weights among BLA and CMA target network links were further analyzed and visualized by BrainNet Viewer (Xia et al., 2013).

## 3. Results

### 3.1 Time-varying intrinsic functional connectivity patterns of the BLA and CMA

We first conducted separate seed-based functional connectivity analyses for the BLA and CMA seeds (**Fig. 1A**). As shown in **Fig. 1B**, we observed a set of widespread brain regions whose spontaneous activity is significantly correlated with the BLA and CMA seeds, with overlapping regions in the insula and the medial frontal cortex as well as anterior and middle portions of cingulate cortex (**Fig. S1**). To further examine the differences between BLA and CMA target networks, we conducted whole-brain 2-by-2 ANOVA with subregions (BLA vs. CMA) and hemispheres (left vs. right) as within-subject factors to identify significant clusters exhibiting the main effects of BLA vs. CMA (**Fig. 1C**). Critically, we observed prominently dissociable patterns of BLA and CMA target connections, involving widespread cortical and subcortical structures. Together with findings from previous studies (Roy et al., 2009; Etkin et al., 2009; Qin et al., 2012), we selected 20 ROIs from five target networks of our major interest in a set of widely distributed network (see Methods), including 1) limbic and paralimbic structures, 2) uni- and polymodal association cortex, 3) subcortical structures, 4) Cerebellum and 5) prefrontal cortex. The rationale of selection for five target networks of interest is depicted in **Fig. 1D** and **Table. S1**.

We then examined time-varying intrinsic functional connectivity properties of 20 target ROIs within five major networks associated with the BLA and CMA, by implementing a sliding window approach in conjunction with K-means clustering methods (**Fig. 1E**). The optimal number of clusters were determined by Silhouette coefficient (Rousseeuw, 1987) and CalinskiHarabasz criterion (Caliński and Harabasz, 1974), with higher value indicating larger cohesion and less dispersity. This analysis revealed the highest scores for both Silhouett coefficient and CalinskiHarabasz criteria at cluster numbers set as 2 (**Fig. S2**). To ensure the stability of time-varying intrinsic functional connectivity and the reliability of clustering, we additionally computed the two metrics using a set of window length (α) and steps (β) varying from 30 to 50 TRs and 1 to 3 TRs respectively. We consistently found optimal states at 2 (**Fig. S2**). We also tested whether BLA and CMA seeds in the two hemispheres showed distinct intrinsic connectivity with target regions or not. Again, we observed a highly similar pattern of results even separating seeds of BLA and CMA for each hemisphere (**Fig. S3**). Together, these results revealed consistent patterns of outcomes, with two clearly dissociable states of time-varying BLA and CMA intrinsic connectivity patterns (**Fig. 2A**).

**Fig. 2.**
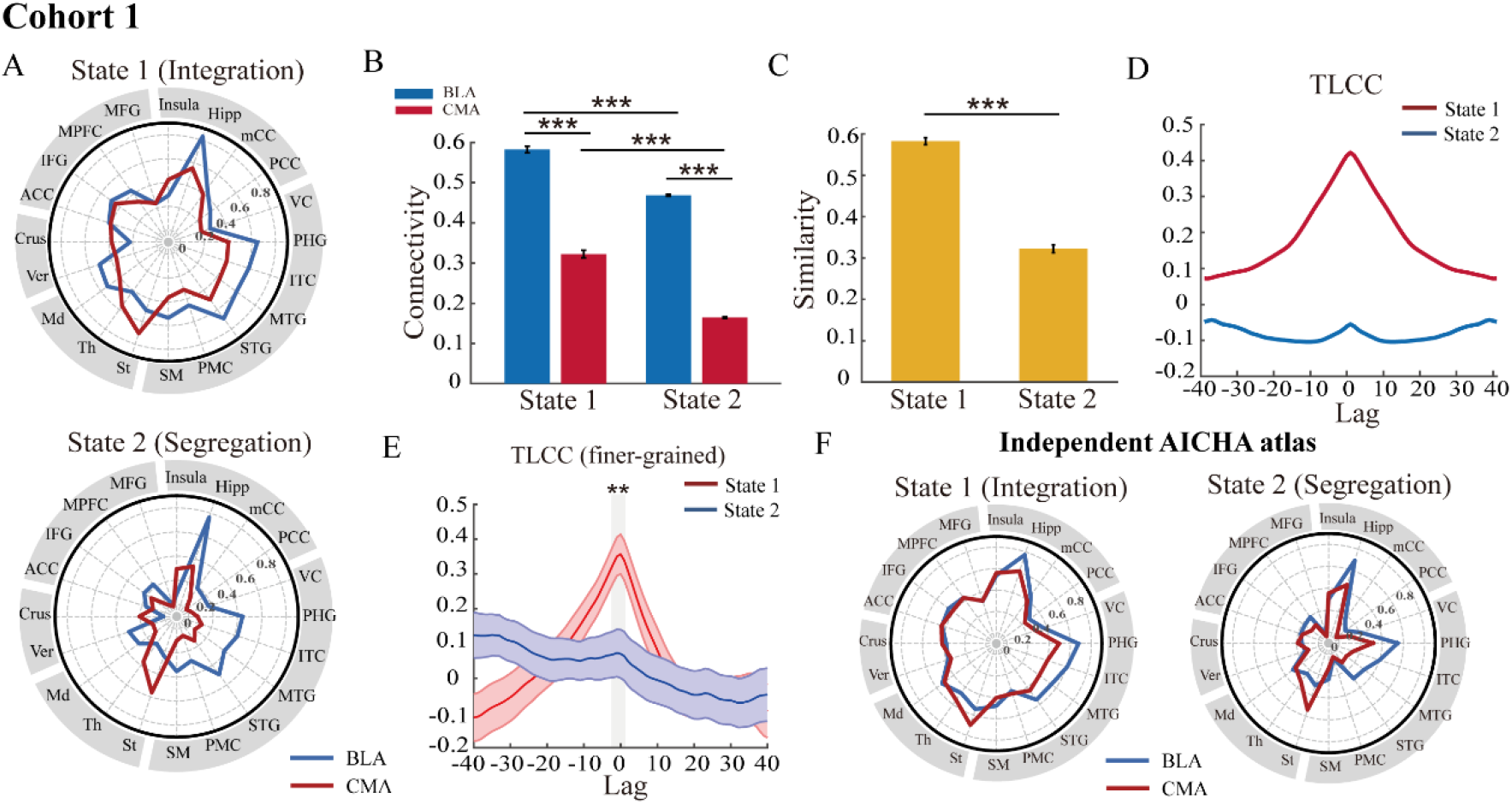
Time-varying integration and segregation states related to distinct spatial-temporal patterns for BLA and CMA connectivity networks (Cohort 1). (**A**) Polar plots depict two distinct states, with generally stronger connectivity strength of both BLA (blue) and CMA (red) with target ROIs for integration state, while significantly weaker yet dissociable connectivity between BLA and CMA seeds. (**B**) Bar graphs show significant difference in averaged connectivity strength (Fisher-z transformed) between BLA-(blue) and CMA-based target regions (red) within and between each state. (**C**) Bar graphs shows averaged correlation of BLA- and CMA-target connectivity networks (similarity) within State 1 and State 2. (**D**) The curves depict time-lagged cross-correlations (TLCC) between BLA and CMA target networks for State 1 and State 2 respectively. (**E**) The curves depict averaged TLCC (finer-grained) along with standard error of the mean for State 1 and State 2 separately. Gray area marks the significant difference between state 1 and state 2 at lag of zero. (**F**) Reproducible dynamic states using a set of independently defined ROIs from the AICHA atlas. Notes: \**P* < 0.05; \*\**P* < 0.01; \*\*\**P* < 0.001, FDR corrected.

### 3.2 Integration and segregation states linking to distinct tempo-spatial BLA and CMA connectivity patterns

Next, we examined tempo-spatial connectivity patterns of the BLA and CMA seeds with target ROIs. As shown in **Fig. 2A**, K-means clustering robustly detected two distinct states, with one state (denoted as State 1) exhibiting generally stronger connectivity strength of both BLA and CMA with target ROIs compared with the other state (denoted as State 2) exhibiting significantly weaker yet dissociable connectivity between BLA and CMA seeds. We then calculated occupancy rate of these two states, and found two states underwent highly time-varying or dynamic changes (**Fig. S4**). To compare intrinsic functional strength of BLA and CMA-target networks within and between State 1 and State 2 (**Fig. 2B)**, we conducted separate independent t tests for averaged functional connectivity (Fisher-z transformed) across windows. Results revealed significantly higher BLA-based connectivity strength than that of CMA, in both State 1 [t_(4193)_ = 46.51, two tailed *p* < 0.001] and State 2[t_(4037)_ = 50.10, two tailed *p* < 0.001]. This pattern of differences is consistent with previous findings in adults (Roy et al., 2009; Qin et al., 2012). More interestingly, we found both BLA-associated and CMA-associated connectivity in State 1 are significantly higher than that in State 2 [separate independent t tests, BLA: t_(8230)_ = 77.82, two tailed *p* < 0.001; CMA: t_(8230)_ = 107.16, two tailed *p* < 0.001]. Additionally, these results could be replicated from edge-by-edge comparisons (**Table. S2, S3)**. Overall, these results indicate distinct difference between the two states, as one showing higher amygdala subregional connectivity with widespread target regions in cortical areas and subcortical structures than the other.

We further investigate spatial similarity and temporal coherence between BLA and CMA target networks within each state. We first calculated spatial similarity between BLA and CMA target networks for the two states, measured by Pearson correlation values (Fisher’s r-to-z transformed) between BLA and CMA-target connectivity across windows (**Fig. 2C**). This analysis revealed significantly higher spatial similarity between BLA and CMA target networks in State 1 (r = 0.58), when compared to that in State 2 [r = 0.32, two sample t-test, t_(8230)_ = 20.47, two tailed *p* < 0.001]. We then calculated time-lagged cross-correlation (TLCC) between BLA and CMA target networks (see Methods). We consistently found higher correlation around lag of zero (**Fig. 2D, Fig. S5)**. Further test on TLCC (finer-grained) at lag of zero revealed significantly higher correlation between BLA and CMA target networks [separate independent t test, t_(65)_ > 3.06, two tailed *p* = 0.003 corrected] (**Fig. 2E**). Notably, all p-values above are FDR corrected for multiple comparisons. We also tested the robustness of results by implementing another finer-grained AICHA atlas (see Methods) and consistently found two distinct states with highly similar tempo-spatial patterns above (**Fig. 2F, Fig. S6**).

We further tested whether the two distinct states are reproducible in an independent cohort of participants (**Cohort 2**) using the same target ROI masks derived from Cohort 1. Again, we found highly similar two distinct states, with one state exhibiting integrated patterns between BLA and CMA-target ROIs compared with the other state (**Fig. 3A**) To be specific, we observed significantly higher BLA and CMA intrinsic connectivity strength in State 1 than that in State 2 [separate independent t tests, BLA: t_(7435)_ = 100.64, two tailed p < 0.001; CMA: t_(7435)_ = 90.42, two tailed p < 0.001] (**Fig. 3B**). Further independent t tests for averaged intrinsic connectivity across windows (Fisher-z transformed) revealed significantly higher BLA-based connectivity strength than that of CMA, in both integration state [t_(2924)_ = 44.19, two tailed *p* < 0.001] and segregation state [t_(4511)_ = 41.21, two tailed *p* < 0.001] (**Fig. 3B**). We also found significantly higher spatial similarity [two sample t-test, t_(7435)_ = 16.17, two tailed *p* < 0.001] (**Fig. 3C)**, and higher TLCC around lag of zero (**Fig. 3D, Fig. S7**). Consistently, test on TLCC (finer-grained) at lag of zero revealed significantly higher correlation between BLA and CMA target networks [separate independent t test, t_(59)_ > 3.44, two tailed *p* = 0.001 corrected] (**Fig. 3E**). Notably, these results were reproducible by using another finer-grained AICHA atlas in both Cohorts 1 and 2 (**Fig. 3F, Fig. S8**).

**Fig. 3.**
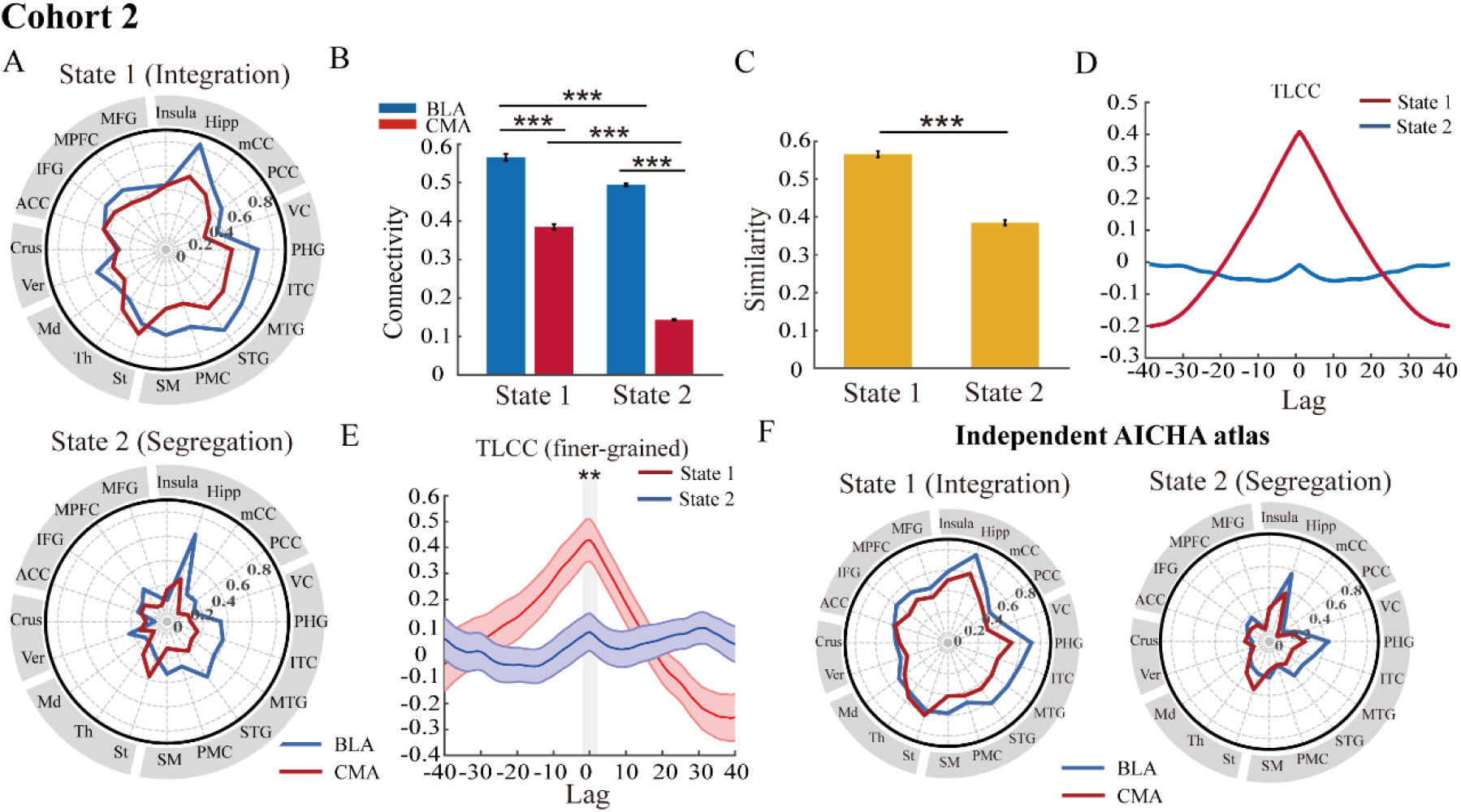
Reproducible results for integration and segregation states with distinct spatial-temporal patterns for BLA and CMA intrinsic connectivity networks in Cohort 2. (**A**) Polar plots depict two distinct states, with generally stronger connectivity strength of both BLA (blue) and CMA (red) with target ROIs for integration state, while significantly weaker yet dissociable connectivity between BLA and CMA seeds. (**B**) Bar graphs show significant difference in averaged connectivity strength (Fisher-z transformed) between BLA-(blue) and CMA-based target regions (red) within and between each state. (**C**) Bar graphs shows averaged correlation of BLA- and CMA-target connectivity networks (similarity) within State 1 and State 2. (**D**) The curves depict time-lagged cross-correlations (TLCC) between BLA and CMA target networks for State 1 and State 2 respectively. (**E**) The curves depict averaged TLCC (finer-grained) along with standard error of the mean for State 1 and State 2 separately. Gray area marks the significant difference between state 1 and state 2 at lag of zero. (**F**) Reproducible dynamic states using a set of independently defined ROIs from the AICHA atlas. Notes: \**P* < 0.05; \*\**P* < 0.01; \*\*\**P* < 0.001, FDR corrected.

Together, by implementing sliding-window approach with K-means clustering method, we found a clear dissociation of tempo-spatial evolution patterns for BLA- and CMA-based intrinsic connectivity within two states. Based on a generally higher BLA- and CMA-based intrinsic connectivity with all of target networks for State 1 and relatively weaker yet dissociable connectivity patterns for State 2, we therefore denote them as integration and segregation states respectively for the present study.

### 3.3 Time-varying states of BLA and CMA connectivity predict spontaneous autonomic arousal

After detecting two dynamic states of amygdala subregions networks, we further investigated whether they could be robustly linked to spontaneous autonomic arousal, measured by tonic-like component of skin conductance level (SCL). We implemented **Cohort 2** (N=37) with concurrent recordings of spontaneous SCL accompanying with above detected dynamic states of BLA and CMA target networks. We first preprocessed SCL data (see Methods) and excluded 7 participants based on the criteria stated in the Methods above and **Fig. S9**. To test the removing out effect, we also conducted additional analysis by including all of participants. We first conducted two independent sample t tests and revealed significantly higher SCL in integration than that in segregation state [separate t-test, before exclusion: t_(7435)_=3.84, *p* < 0.001; after exclusion: t_(6028)_=3.23, *p* = 0.0012] (**Fig. 4A)**.

**Fig. 4.**
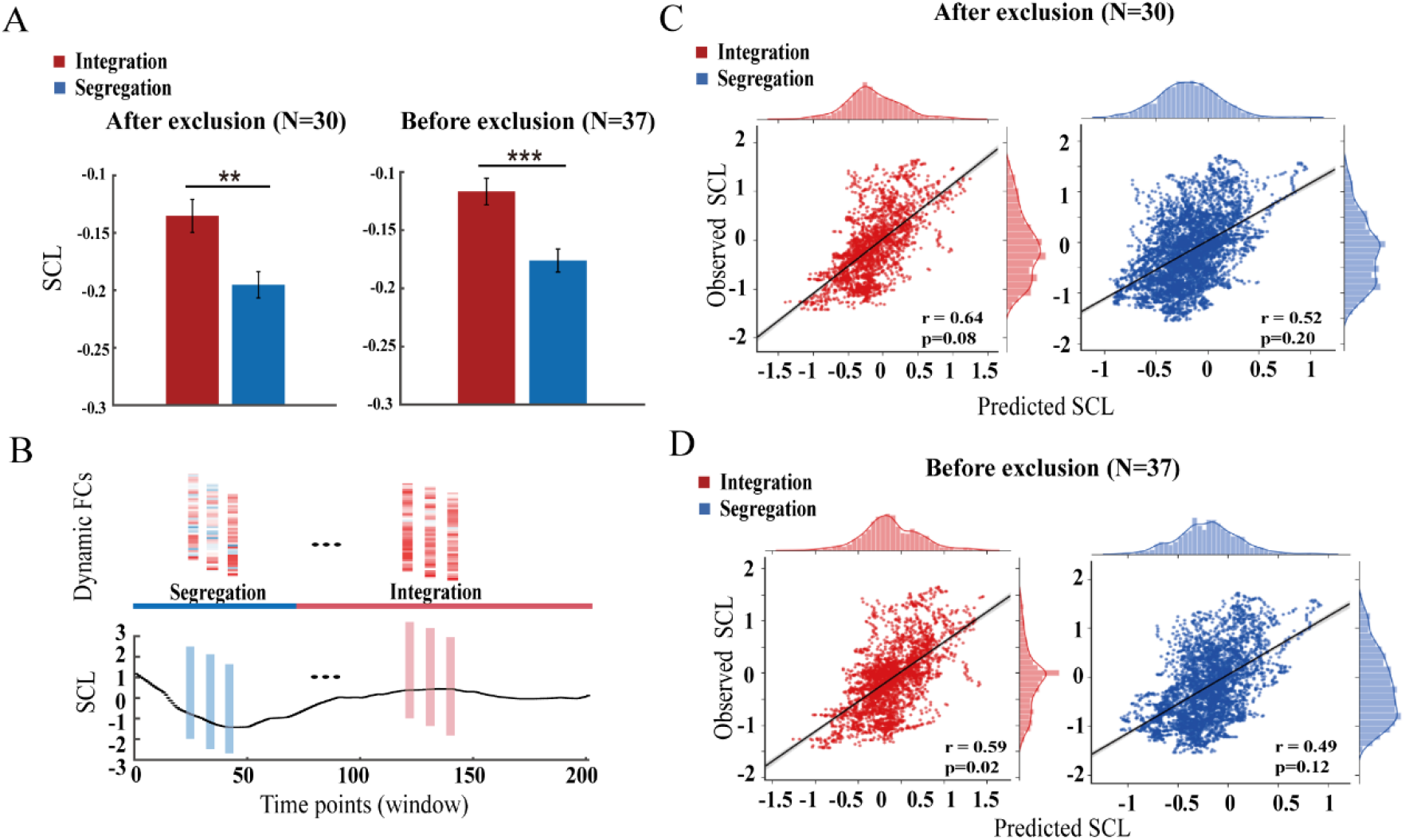
Time-varying integration and segregation states linking to spontaneous skin conductance level (SCL). (**A**) Bar graphs depict significant differences in skin conductance levels between integration and segregation states before and after exclusion of 7 participants. (**B**) An example of dynamic functional connectivity for integration and segregation states as a function of spontaneous SCL fluctuations from one participant. (**C**) Scatter plots show correlations between predicted and observed SCL values for two states with exclusion of 7 participants (N=30). Predicted SCL value are derived from Elastic-net regression for integration and segregation states respectively. Shaded areas represent 95% confidence interval. (**D**) Same plot with **C** but without exclusion of 7 subjects (N=37). Notes: \**P* < 0.05; \*\**P* < 0.01; \*\*\**P* < 0.001.

Furthermore, we implemented Elastic-net regression (Zou et al., 2005; Cui et al., 2018) to examine whether time-varying functional connectivity networks of BLA and CMA seeds could be predictive of ongoing SCL fluctuations. Two Elastic-net regression models were trained for BLA and CMA-based intrinsic connectivity data for time-varying windows within integration and segregation states separately. Each window-wise connectivity from integration and segregation states is considered as a function of concurrent recordings of window-wise SCL date (**Fig. 4B**). As mentioned in Methods, we employed grid-search for determining optimal parameter set (*α, λ*) (**Fig. S10**). Then, to determine the significance level for predictive accuracy in Elastic-net models, we employed a stringent permutation test to control autocorrelations inherent in functional connectivity metrics and SCL time series (see Methods). Outcomes from the permutation tests revealed prominent correlations between the predicted and observed spontaneous SCL fluctuations within integration state [before exclusion: *r* = 0.59, permutated *p* = 0.02; after exclusion: *r* = 0.64, permutated *p* = 0.08]. However, we did not observe this association for segregation state [before and after exclusion: permuted *p* = 0.12 and *p* = 0.20 respectively] (**Fig. 4C, 4D**). When implementing conventional permutations, associations of time-varying BLA/CMA connectivity with spontaneous SCL fluctuations were both significant in integration and segregation states with and without exclusion (all permutated *p <* 0.001). To further examine the differences in predictive accuracy under integration and segregation states, we implemented nonparametric bootstrap approaches. Independent t test revealed a significantly higher predictive accuracy for SCLs in the integration than segregation state [before exclusion: t_(9998)_ = 328.48, two tailed *p* < 0.001; after exclusion: t_(9998)_ = 259.19, two tailed *p* < 0.001], indicating higher correlations between time-varying BLA/CMA intrinsic connectivity and SCL fluctuations in integration state, compared with segregation state. Together, these results indicate higher predictive correlation between time-varying BLA/CMA intrinsic connectivity and SCL fluctuations for integration than segregation states.

### 3.4 Distinct BLA and CMA target network configurations predict autonomic arousal

We further investigated prediction weights of each connectivity link (or feature) derived from Elastic-net regressions for integration and segregation states respectively. Analyses of BLA and CMA-based target network configurations revealed a dissociation of prediction weights in multiple distributed brain regions between the integration and segregation states (**Fig. 5A, Table. S4, S5**). We then computed the relative ratio of total prediction weights for BLA and CMA-target networks separately. Interestingly, we observed a relatively CMA-dominated contribution in the integration state and a BLA-dominated contribution in the segregation state (**Fig. 5B**). This pattern of results remains stable when accounting for the most contributed connections under different thresholds (**Fig. S11**). We further explored the spatial distribution for most predictive connections by visualizing the top 25% BLA- and CMA-based target connections in integration and segregation states separately (**Fig. 5C**). For the integration state, we observed prominent target connections involving the sensory cortex (SM, STG, ITC), limbic structures (Insula, mCC) and subcortical structures (thalamus). For the segregation state, we observed the top 25% connections in prefrontal systems (i.e., MPFC, MFG, IFG) and (para)hippocampus regions as well as crus regions in cerebellum. Again, such dissociation between integration and segregation states is reproducible even without exclusion of 7 participants (**Fig. S12**). Together, these results indicate distinct BLA and CMA target network configurations in integration and segregation states that predict spontaneous SCL levels.

**Fig. 5.**
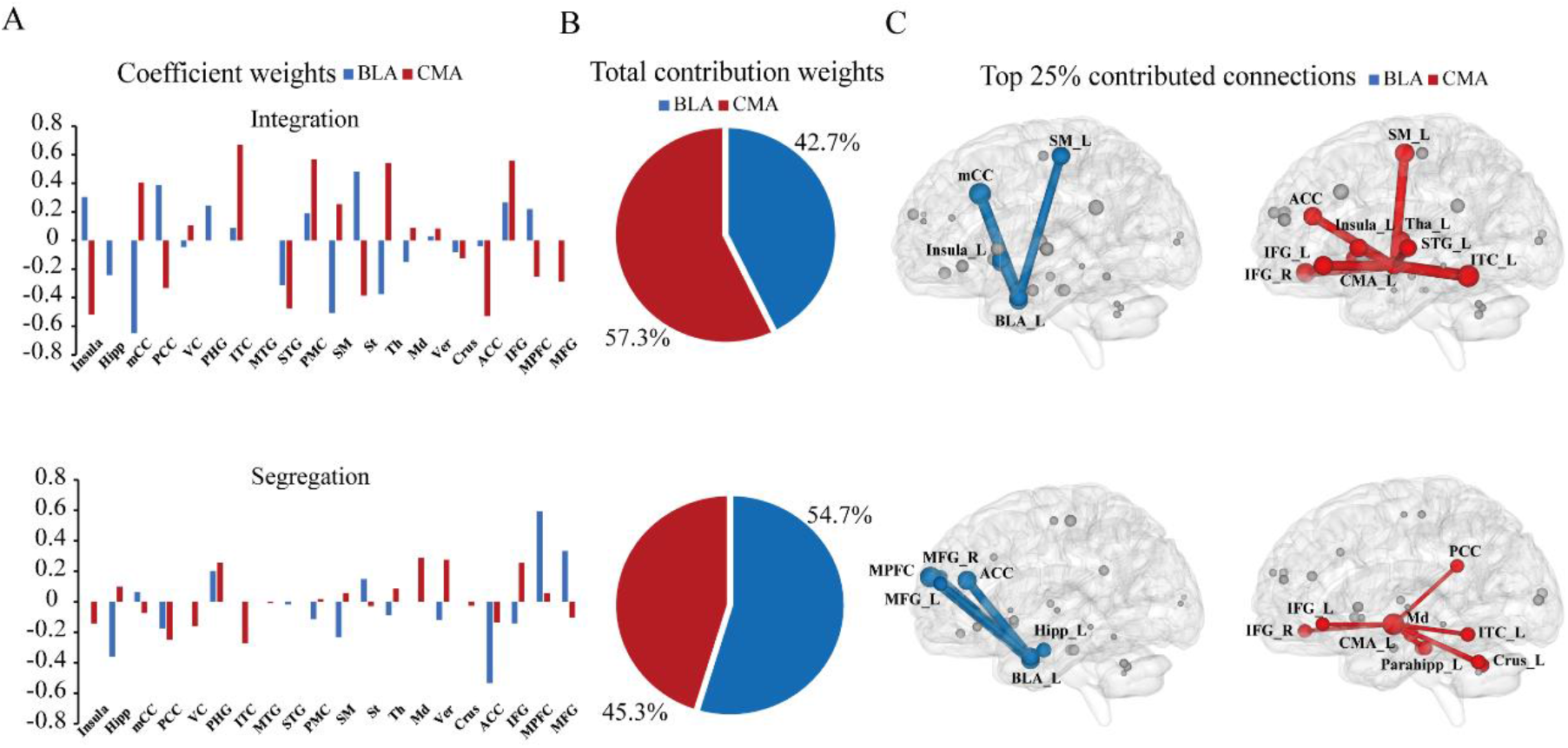
Prediction weights and spatial distributions of BLA and CMA target network configurations linking to spontaneous SCL fluctuations (N=30). **(A)** Bar graphs depict coefficients derived from Elastic-net model, for integration (top) and segregation (bottom) state separately. (**B**) Pie charts represent relative ratio of total prediction weights between BLA- and CMA-target connections in Elastic-net model, for integration (top) and segregation (bottom) state separately. (**C**) Top 25% predictive connections of BLA- (blue) and CMA-based (red) target networks in integration (top) and segregation (bottom) states are superimposed onto the glass brain in a sagittal view separately, with connections in the left hemisphere for visualization purpose only. Size of nodes and corresponding connections represent the relative prediction weights of BLA- and CMA-target links derived from Elastic-net regression model separately.

## 4. Discussion

In this study, we investigated the dynamical organization of intrinsic functional connectivity networks associated with the amygdala nuclei and its contributions to autonomic arousal in humans. By leveraging machine learning and clustering approaches, we identified two distinct states of time-varying BLA/CMA intrinsic connectivity networks: an integration state exhibits generally stronger connectivity among BLA- and CMA-based target networks, whereas a segregation state exhibits relatively weaker yet dissociable BLA-than CMA-based connectivity with cortical regions. Furthermore, we found a clear dissociation of spatio-temporal patterns for BLA- and CMA-based intrinsic connectivity networks within integration and segregation states. Critically, such time-varying dissociable states of BLA- and CMA-based target networks are linked to spontaneous SCL fluctuations – a measure of autonomic arousal, with significantly higher SCL and higher predictive accuracy in the integration as compared to the segregation state. Such time-varying integration and segregation states with distinct BLA- and CMA-based network configurations contribute to predicting spontaneous fluctuations of SCL.

One of our major findings is to identify two distinct states of time-varying intrinsic functional connectivity patterns for BLA and CMA target networks by implementing sliding-window and K-mean clustering methods. Specifically, the integration state exhibits generally stronger connectivity with almost all of target ROIs and substantially overlapping between BLA and CMA target networks, as compared with the segregation state. Unlike the integration state, however, the segregation state exhibits relatively weaker yet clearly functional dissociation between BLA and CMA target networks. This segregation pattern partially is in line with previous findings on functional segregation of BLA and CMA target networks using the conventional static averaging (static-like) approaches (Etkin et al., 2009; Qin et al., 2012). Building on the neuroanatomical models in rodents, with the amygdalar nuclei via unique connections to form dedicated circuits in support of distinct affective functions (LeDoux, 2000; LeDoux, 2007), our observed time-varying integration and segregation states provide prominent evidence to highlight a dynamic functional reorganization of BLA- and CMA target networks over time in humans. Moreover, BLA and CMA target networks in integration state showed a higher spatio-temporal coupling phenomenon compared with segregation state, suggesting an integrated property for integration state. Such highly coupling patterns between BLA and CMA target networks, in a spatio-temporal manner, are not reported by the conventional scan-length averaging functional connectivity approaches. Notably, such dynamic integration and segregation pattern is reproducible by another independent cohort of participants and by using a totally independent AICHA atlas in both cohorts, suggesting the reproducibility and robustness of our current findings. Together, these two dynamic states provide novel evidence to suggest that BLA and CMA intrinsic functional circuits do not remain constant, rather undergo spontaneously fluctuated over time. This extends previous findings on intrinsic functional organization of the amygdalar nuclei derived from conventional (static-like) approaches and animal models (Roy et al.,2009; Qin et al., 2012).

A second major finding of our study is that SCL during the integration state is significantly higher than segregation state. Since SCL is a reliable measure of autonomic arousal (Critchley et al., 2010; Boucsein, 2012), it is thus conceivable that our observed associations may reflect a link between time-varying functional connectivity of the amygdala nuclei and autonomic (physiological) arousal. Notably, the window-averaging SCL in our present study most likely reflects a tonic-like component of autonomic arousal rather than external stimulus-induced SCR in previous studies (Patterson et al., 2002; Bach et al., 2009, 2010). In other words, our observed relatively slow fluctuations of SCL may represent spontaneity of one’s internal autonomic arousal to some extent, which is reminiscent of findings from one recent study (Baczkowski et al., 2017). And the BLA- and CMA-based functional circuits and their associated intrinsic networks could play an essential role in modulating such internal state. From a systems level, converging evidence from human functional neuroimaging suggests large-scale configurations in functional brain networks detected by BOLD-fMRI signals are closely linked to changes of autonomic arousal due to the activation of (non)adrenergic systems (Chang et al., 2009; Phelps et al., 2005; Hermans et al., 2011; Kim et al., 2017). Animal studies have also demonstrated that LC releasing norepinephrine (Aston-Jones & Cohen, 2005) acts on corresponding receptors in widely distributed brain regions especially the amygdala and related networks, thereby prompting rapid increases in neuronal excitability and functional coupling among these disparate regions (Arnsten, 1998).

Moreover, we found that time-varying amygdala subregional intrinsic connectivity patterns in the integration state were predictive of spontaneous fluctuations of SCL, with significantly higher predictive accuracy in the integration than segregation state. These may reflect increased functional coordination of BLA and CMA-modulated large-scale neural networks, accompanying with highly associated autonomic arousal measured by pupillometry data in humans (Murphy et al., 2011). Such association has been observed in animal models. For example, acute stress can trigger a widespread cascade of neurochemical changes that rapidly prompt neuronal excitability of large-scale brain networks into a tonically sensitive mode (Joëls & Baram, 2009; Stujenske & Likhtik, 2017). Thus, it is possible that there are multiple neuromodulatory systems contributing to our observed stronger yet overlapping pattern between BLA and CMA target networks in the integration state. In other words, the integration state may result from endogenous autonomic and hormonal stimulations promoting the excitability of neuronal ensembles in the amygdalar nuclei and related functional circuits, thereby promoting the adapative flexibility of the brain’s internal affective or homeostatic states to meet ever-changing environmental needs. Indeed, evidence from recent studies have suggested that integrated states may enable faster and more effective performance on a cognitive task, most likely through ascending neuromodulatory systems to regulate the transition between multiple modes of brain function (Shine et al., 2016; Shine et al., 2019).

A third important finding of our study is that distinct features (or connections) from time-varying states of BLA and CMA target networks contribute to predicting spontaneous autonomic arousal as evidenced by SCL fluctuations. Specifically, a CMA-dominated pattern relating to predicting autonomic arousal in integration state was observed, compared with a BLA-dominated pattern in segregation state. Such divergence between two dynamic states may exhibit distinct arousing modes, with one owning preference of CMA-engagement phenomenon (integration state) and the other exhibiting preference of a BLA-engagement pattern (segregation state). Additionally, such dynamical phenomenon is to some extent in line with newly emerging evidence from recent animal research, which demonstrates that amygdala ensembles can dynamically represent internal affective or homeostatic states (Janak et al., 2015; Beyeler et al.,2018). Specifically, the dynamic assembly of two major populations of neurons in the BLA plays a role in regulating animal’s internal autonomic or physiological states through several output pathways to larger brain networks (Salzman et al., 2010; Zhang et al., 2018; Grundemann et al., 2019). However, given the fact that neuroanatomical and neurochemical properties of the amygdala nuclei in rodents may differ from that in humans (Pabba et al., 2013), future studies are required to elucidate the neurobiological mechanisms of how the BLA- and CMA-based functional circuits regulate spontaneous autonomic arousal in humans.

Moreover, the divergence between two dynamic states is further supported by their distinct spatial distributions for the BLA- and CMA-based intrinsic connectivity among widespread brain regions, including portions of prefrontal cortex, cingulate cortex, (para)hippocampus, sensory-motor cortex as well as cerebellum regions. These dissociable patterns may reflect a dynamic nature of the amygdalar nuclei (BLA and CMA) in converging information from multiple brain systems closely related to affective, homeostatic and cognitive states (Craig, 2002; Critchley, 2005; Pollatos et al., 2007). Given the fact that dynamic states are robustly associated with SCL, we assume that functional coordination of these regions could further reflect an increase in neuronal excitability and functional connectivity within the salience network that is responsible for hypervigilant, anxious and/or stressful states (Hermans et al., 2011; McMenamin et al., 2014). From a psychopathological perspective, both hyper-activation and hyper-connectivity in the amygdala nuclei and associated functional circuits have been linked to internalizing symptoms and abnormalities in affective functions and autonomic responses including somatization symptoms and anxiety (Bishop, 2007; Hermans et al., 2011). Thus, the dynamical organization of integration and segregation states within the amygdala nuclei and their intrinsic networks could be relevant to develop the objective biomarkers linking to internalizing symptoms and autonomic arousal in affective disorders. In addition, the amygdala is recognized to play a critical role in affective information processing likely through its functional coupling with regions in polymodal association areas and the prefrontal cortex in emotion perception, appraisal and regulation (Lim et al., 2009). Thus, it is likely to speculate that our observed BLA and CMA target networks in the integration and segregation states may reflect internal physiological responses with different responsive modes of arousal involved in a variety of cognitive and affective functions that can’t be detected by the conventional methods. Furthermore, the dynamic transitions between these two distinct states may together build self-organized dynamics modulating temporally rapid autonomic arousal in a systematic level.

The dynamical (re)organization of large-scale brain networks holds the promise to enable the flexibility of brain functioning and rapid adaptation in face of ever-changing environmental demands (Deco et al.,2011; McMenamin et al., 2014; Braun et al., 2015). To our knowledge, no studies to date have directly investigated the dynamical organization of amygdala subregional functional circuits and networks. Our findings thus extend the existing knowledge on functional organization of emotion-related brain circuitry which is theorized to play a critical role in regulating one’s internal autonomic (and more broadly homeostatic) states. This line of research has its potential to extend into investigating function and dysfunction of emotion-related brain circuitry in affective disorders including anxiety and depression. The mainstay of previous studies using the conventional approaches have demonstrated emotion-related brain dysconnectivity in patients with affective disorders, with hyper- and/or hypo-connectivity patterns in limbic, paralimbic and prefrontal systems critical for vigilance, rapid detection and memories for threatening stimuli, emotion regulation and expression (Adolphs et al., 1994; Clark, & Beck, 2010; Sturm et al., 2003). Thus, it would be highly relevant to investigate how dysregulated brain dynamics as well as functional circuits linked to the amygdalar nuclei in particular underlie the core symptoms of affective disorders. Understanding the dynamical organization of emotion-related brain circuitry represents an important step toward developing brain-inspired biomarkers of affective and psychosomatic disorders on a circuit-level (Insel et al., 2010). Future studies are required to delineate how the dynamical organization of the amygdala nuclei with broader networks regulates internal affective or autonomic responses in both healthy and psychiatric conditions.

Our findings should be considered in light of several limitations. First, the sliding window and machine learning methods we implemented here rely on ad hoc procedures to determine parameters like window length or clustering numbers. More precise and unbiased approaches are needed in future studies. Second, there is a relatively small sample size for SCL data in our present study, which may limit its generalizability to other populations. Third, our study highlights the association of time-varying states of BLA- and CMA-based intrinsic functional connectivity networks with spontaneous autonomic arousal, and future studies are needed to shed light on the mechanistic understanding of how amygdala subregions dynamically interact through specific projections to modulate autonomic arousal and internal states in humans.

**In conclusion**, our study demonstrates dynamic integration and segregation of the emotion-related brain networks that are linked to different internal autonomic states. Our findings provide important implications into understanding the neurobiological mechanisms of how the dynamical organization of emotion-related brain networks regulates internal physiological states, and eventually inform dysregulated brain dynamics in affective disorders and psychopathology.

## Acknowledgements

This work was supported by the National Natural Science Foundation of China (31522028, 81571056, 2014NT15), the Open Research Fund of the State Key Laboratory of Cognitive Neuroscience and Learning (CNLZD1503), the Major Project of National Social Science Foundation (19ZDA363), the Major Project of National Social Science Foundation (20&ZD153), the BNU Interdisciplinary Research Foundation for the First-Year Doctoral Candidates (BNUXKJC1910).

