## Supplemental Table 1-5, Figure 1-11 for "Dynamic integration and segregation of amygdala subregional functional circuits linking to physiological arousal"

### Supplement Materials

#### Supplementary Figures

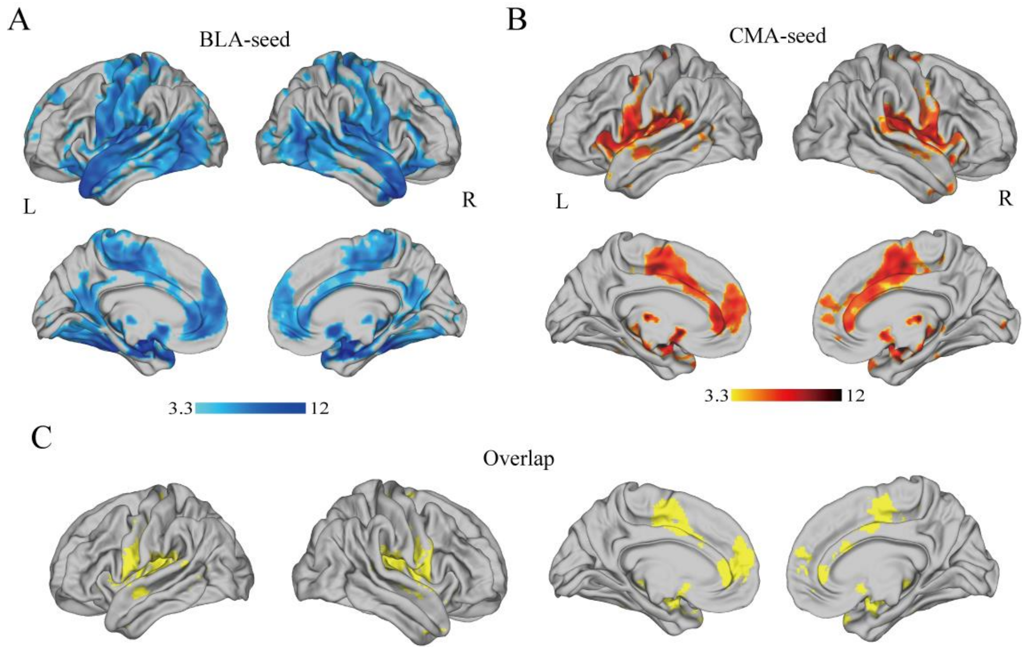


**Fig. S1. BLA- and CMA-seeded intrinsic functional connectivity maps.** (Top) The lateral views of significant clusters derived from intrinsic functional connectivity of the BLA (blue) and CMA (red) seeds. (Bottom) Overlaps between BLA and CMA-target connectivity maps in yellow. Color bar represents connectivity strength. Notes: L, Left; R, Right.

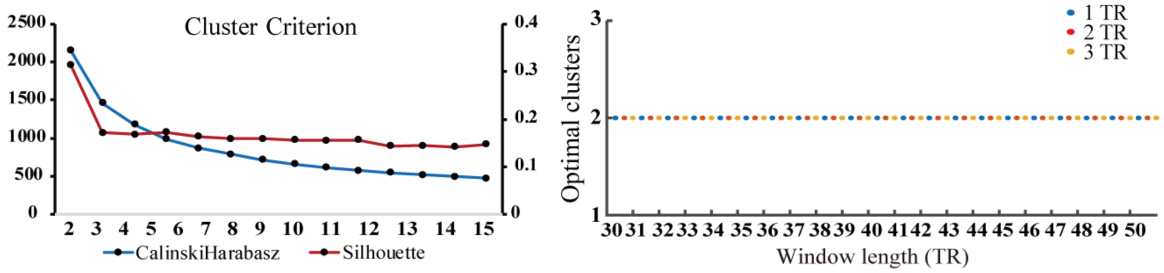


**Fig. S2. Objective criterion for determining optimal clustering numbers.** (Left) Line curves depict the CalinskiHarabasz and Silhouette criteria as a function of cluster number K. (Right) Different combinations of window length (30 TRs to 50 TRs) and step (1 TR to 3 TRs) were selected to test the robustness of optimal clustering numbers.

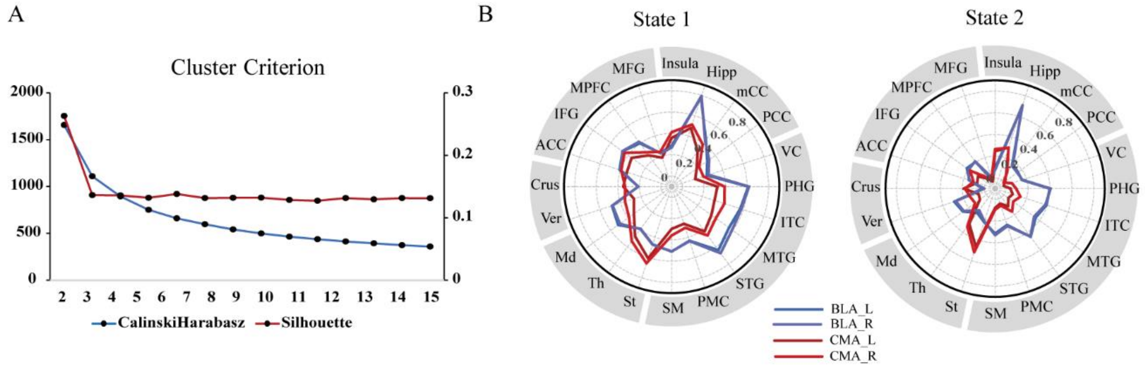


**Fig. S3. Clustering results for BLA and CMA-seeds at each hemisphere separately (Cohort 1).** We computed BLA and CMA-seed target connectivity networks for each hemisphere separately and conducted the same k-means clustering analysis. (**A**) Line curves depict the CalinskiHarabasz and Silhouette criteria as a function of cluster number K, consistently showing optimal clustering numbers at 2. (**B)** Polar plots depict two distinct states with BLA and CMA-seeds target connectivity patterns for each hemisphere, which are similar with those in **Fig. 2A**.

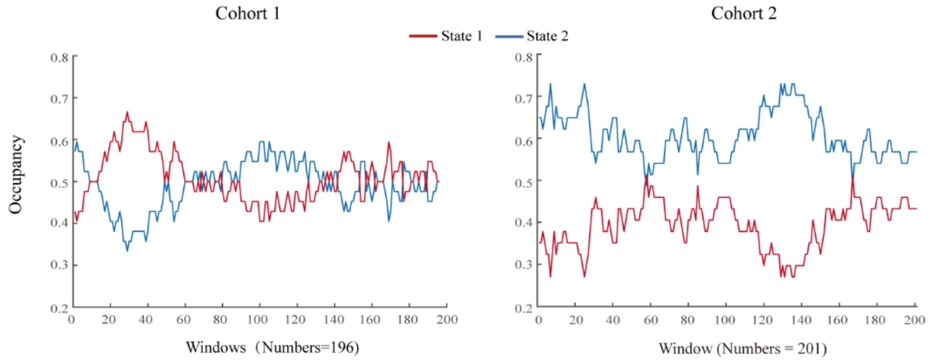


**Fig. S4. State occupancy rate during each window in two cohorts.** The occupancy rate for State 1 (red) and State 2 (blue) across all the participants are calculated by post hoc assignment after K-means clustering and plotted at each window. The x-axes represent total window counts for **Cohort 1** and **Cohort 2** and the y-axes represent occupancy of the two states across participants under each window.

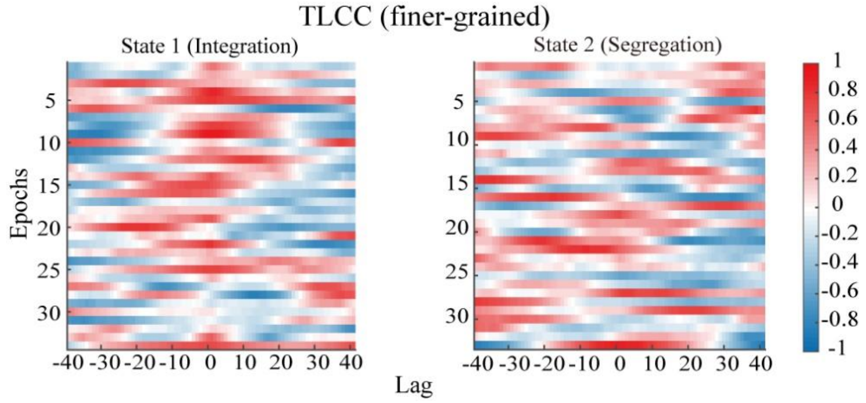


**Fig. S5. Finer-grained TLCC for between BLA and CMA-target connectivity networks for each state (Cohort 1).** (**A**) Heat maps depict finer-grained time-lagged cross-correlation (TLCC) for state 1 and state 2 (see Methods). X axis represent lags and y axis represent each epoch and color represents relative correlation value. Notes**:** Total 34 epochs for the state 1 and 33 epochs for the state 2.

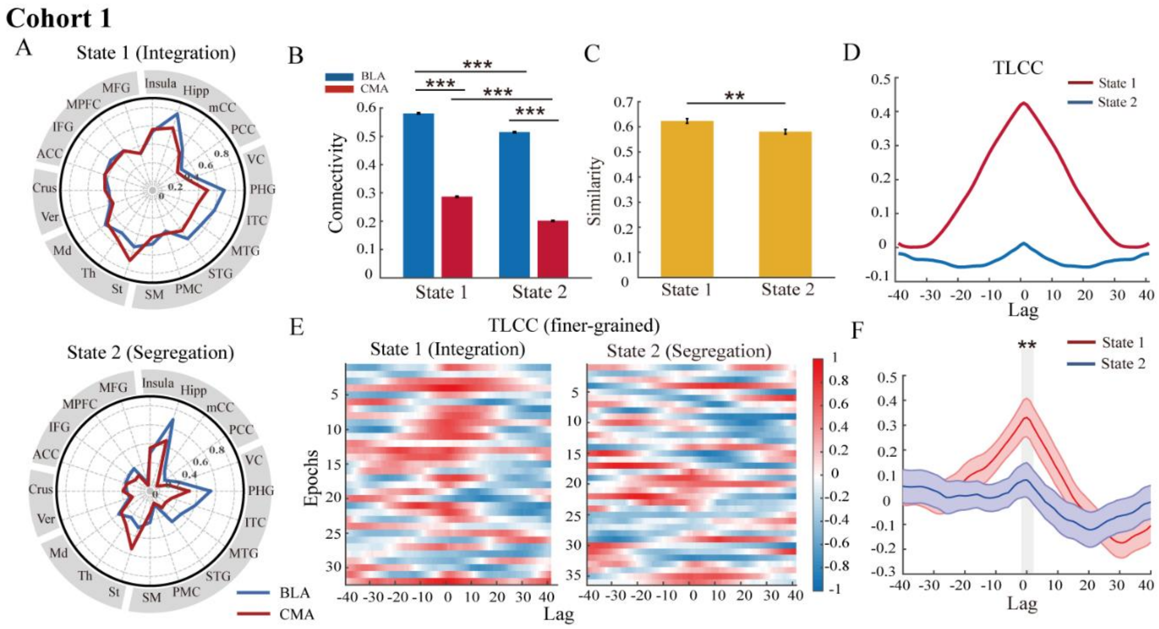


**Fig. S6. Replication results for state analysis using AICHA atlas (Cohort 1).** Polar plots depict two distinct states similar with those in Cohort 1. (**B**) Bar graphs (top) shows significant difference in connectivity strength between BLA- (blue) and CMA-based target regions (red) for each state. (**C**) Bar graphs (bottom) shows averaged correlation of BLA- and CMA-target connectivity networks within integration and segregation state, respectively. (**D**) The curves depict time-lagged cross-correlations between BLA and CMA target networks for state 1 and state 2. (**E**) Heat maps depict TLCC (finer-grained) for State 1 and State 2. The x-axes represent cross-correlation between BLA- and CMA-target networks under different lags. The y axis represents each epoch and color represents correlation value. Total 32 epochs for the state 1 and 36 epochs for the state 2. (**F**) The curves depict averaged TLCC (finer-grained) along with standard error of the mean for State 1 and State 2 separately. Gray area marks the significant difference between state 1 and state 2 at lag of zero. Notes: **P* < 0.05; ***P* < 0.01; ****P* < 0.001.

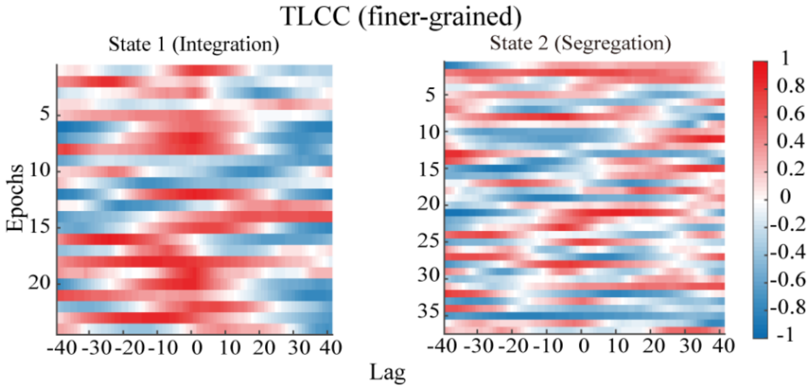


**Fig. S7. Finer-grained TLCC for between BLA and CMA-target connectivity networks for each state (Cohort 2).** (**A**) Heat maps depict finer-grained time-lagged cross-correlation (TLCC) for state 1 and state 2 (see Methods). X axis represent lags and y axis represent each epoch and color represents relative correlation value. Notes: Total 24 epochs for the state 1 and 37 epochs for the state 2.

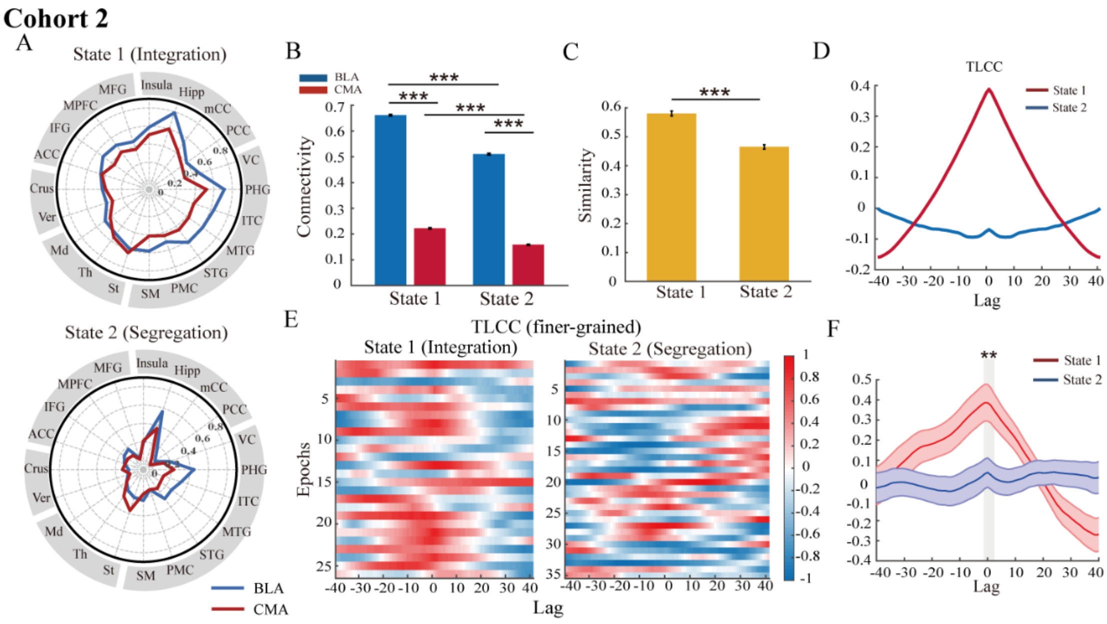


**Fig. S8. Replication results for state analysis using AICHA atlas (Cohort 2).** Polar plots depict two distinct states similar with those in Cohort 1. (**B**) Bar graphs (top) shows significant difference in connectivity strength between BLA- (blue) and CMA-based target regions (red) for each state. (**C**) Bar graphs (bottom) shows averaged correlation of BLA- and CMA-target connectivity networks within integration and segregation state, respectively. (**D**) The curves depict time-lagged cross-correlations between BLA and CMA target networks for state 1 and state 2. (**E**) Heat maps depict TLCC (finer-grained) for State 1 and State 2. The x-axes represent cross correlation between BLA- and CMA-target networks under different lags. The y axis represents each epoch and color represents relative correlation value and color represents relative correlation value. Total 26 epochs for the state 1 and 35 epochs for the state 2. (**F**) The curves depict averaged TLCC (finer-grained) along with standard error of the mean for State 1 and State 2 separately. Gray area marks the significant difference between state 1 and state 2 at lag of zero. Notes: **P* < 0.05; ***P* < 0.01; ****P* < 0.001.

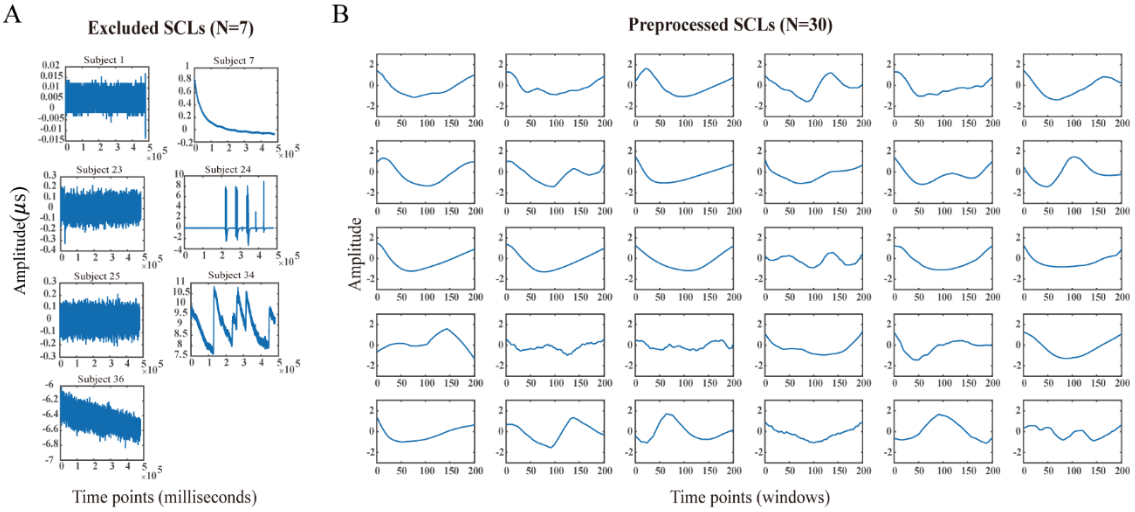


**Fig. S9. Preprocessed skin conductance level in Cohort 2.** (**A**) Before preprocessing, we excluded subject 1,7,23,25,36 for their existence of negative amplitude during recording time and subject 24,34 for relatively large-scale change of amplitude during scanning. x axis represents time points and y axis represent amplitude measured by micro-Siemens. (**B**) Plots show preprocessed skin conductance levels for the remaining subjects (N=30). The y axis in each subfigure represents tonic component drifts and x axis represents time points.

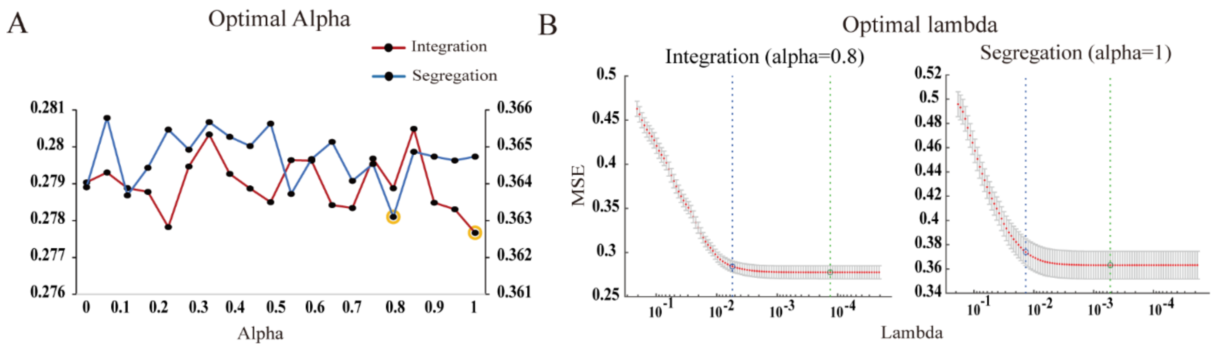


**Fig. S10. Parameter selection procedure for Alpha and Lambda in elastic-net model.** (**A**) For each $\alpha$, we calculated MSE under different $\lambda$ and depicted the minimum MSE at each $\alpha$ value in integration (red line) and segregation (blue line) separately. x axis represents different $\alpha$ while y axis represents values of MSE for integration (red, left) and segregation (blue, right) separately. Yellow circles represent chosen $\alpha$ values for two states separately which has the minimum MSE from all possible parameter settings. (**B**) Plots depict MSE under different $\lambda$ values for integration (left) and segregation (right), after determining the optimal $\alpha$ values. Error bars represents the standard of the MSE computed by 10-fold cross-validation. The green circles indicate the Lambda with one minimum MSE. The blue circles indicate the largest lambda such that the MSE is within one standard error of the minimum MSE.

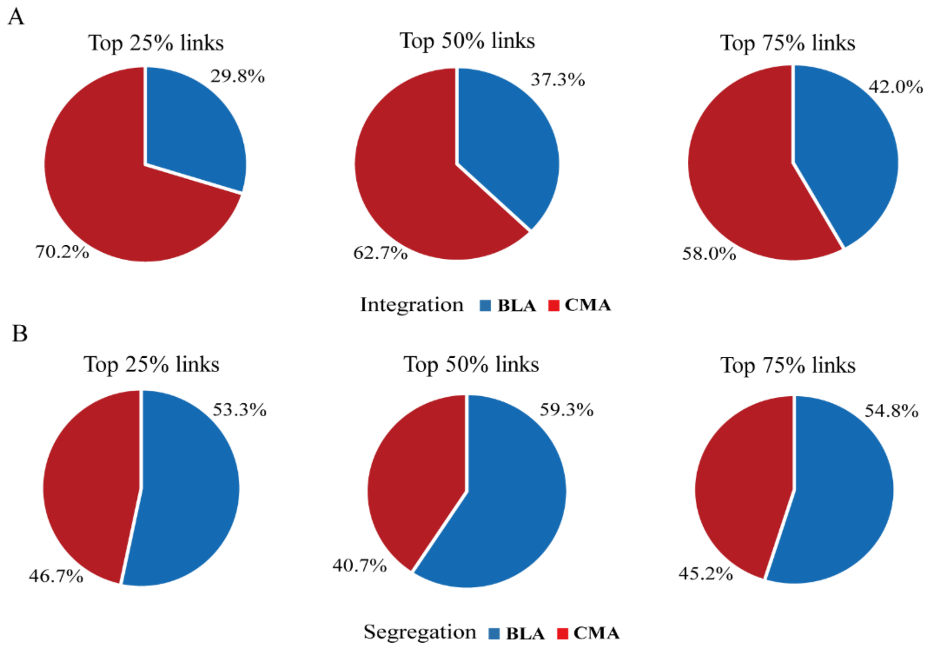


**Fig. S11. Relative contribution ratio between BLA and CMA networks in integration and segregation states under different threshold.** Relative ratio of contribution weights (measured by total contributed weights) between BLA and CMA-target networks are plotted in pie charts for integration (**A**) and segregation (**B**) state separately.

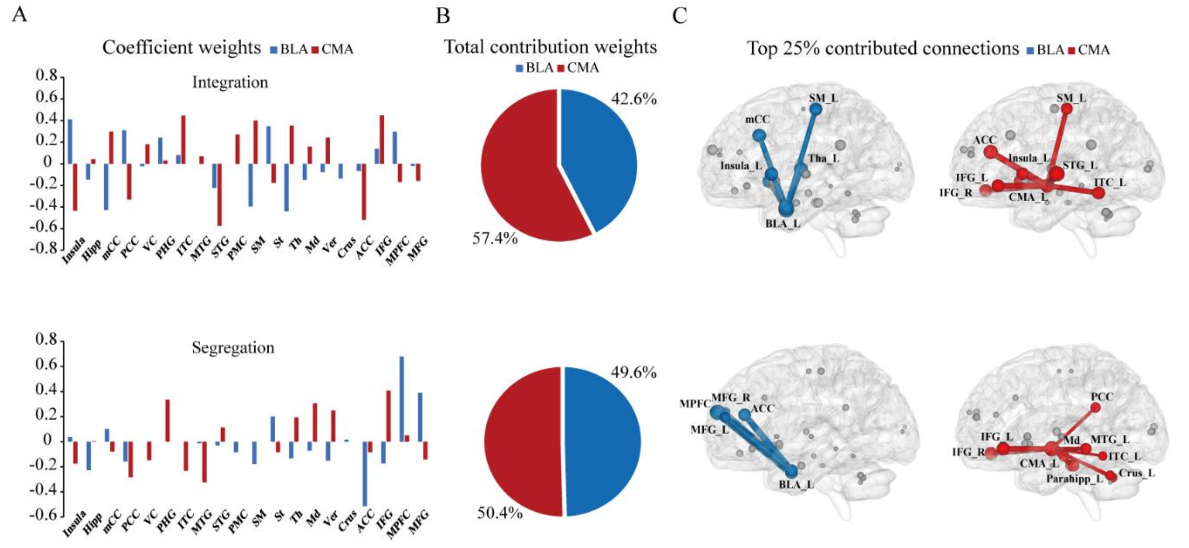

**Fig. S12. Coefficient analysis without exclusion of 7 subjects (N=37). (A)** Bar graphs depict coefficients derived from Elastic-net model, for integration (top) and segregation (bottom) state separately. (**B**) Pie charts represent relative ratio of total prediction weights between BLA- and CMA-target connections in Elastic-net model, for integration (top) and segregation (bottom) state separately. (**C**) Top 25% contributed connections of BLA (blue) and CMA- (red) target networks during the integration (top) and segregation (bottom) are superimposed onto the glass brain in a sagittal view separately, with connections in the left hemisphere for visualization purpose only. Size of nodes and corresponding connections represent the relative contribution weights of BLA- and CMA-target links derived from elastic-net model separately.

#### Supplementary Tables

**Table. S1: The rationale of selection for five target networks of interest**

| Systems or networks | Regions of interest | Affective and cognitive functions |
|---|---|---|
| Limbic and paralimbic structures | Hippocampus, Insula, mCC, PCC | Regulation of arousal, positive and negative affect and motivation |
| Uni- and polymodal association cortex | VC, PHG, ITC, MTG, STG, PMC, SM | Sensory and motor processing for affective information |
| Subcortical structures | Striatum, thalamus and midbrain | Reward and motivation |
| Cerebellum | Crus and vermis/declive regions | Regulating fear related defensive and reflective responses |
| Prefrontal cortex | ACC, MFG, IFG, MPFC | Emotional appraisal and regulation |

Notes: mCC, middle portions of cingulate cortex; PCC, posterior portions of cingulate cortex; VC, visual cortex; PHG, parahippocampal gyrus; ITC, inferior temporal cortex; MTG, middle temporal gyrus; STG, superior temporal gyrus; PMC, premotor cortex; SM, sensorimotor cortex; ACC, anterior cingulate cortex; MFG, middle frontal gyrus; IFG, inferior frontal gyrus; MPFC, medial prefrontal cortex.

**Table. S2. Brain regions showing significant differences in functional connectivity between integration and segregation state, computed separately for CMA- and BLA-seed target networks (Cohort 1)**

| Brain regions | | BLA-network | | CMA-network | |
|---|---|---|---|---|---|
| | | State 1 versus State 2 | | | |
| | | T values | *p* values | T values | *p* values |
| Limbic and paralimbic structures | Insula | 34.17 | *** | 23.74 | *** |
| | Hippocampus | 29.88 | *** | 45.75 | *** |
| | mCC | 49.46 | *** | 59.91 | *** |
| | PCC | 28.26 | *** | 40.65 | *** |
| Uni- and polymodal association cortex | VC | 15.93 | *** | 27.79 | *** |
| | Parahippocampal gyrus | 43.44 | *** | 61.30 | *** |
| | ITC | 40.03 | *** | 57.12 | *** |
| | MTG | 46.61 | *** | 69.91 | *** |
| | STG | 46.08 | *** | 70.80 | *** |
| | PMC | 30.07 | *** | 45.76 | *** |
| | SM | 34.27 | *** | 44.46 | *** |
| Subcortical structures | Striatum | 51.13 | *** | 37.66 | *** |
| | Thalamus | 38.41 | *** | 44.17 | *** |
| | Midbrain | 45.90 | *** | 52.22 | *** |
| Cerebellum | Vermis/declive | 35.83 | *** | 46.59 | *** |
| | Crus | 35.44 | *** | 30.10 | *** |
| Prefrontal cortex | ACC | 44.64 | *** | 56.70 | *** |
| | IFG | 46.38 | *** | 54.80 | *** |
| | MPFC | 39.14 | *** | 56.01 | *** |
| | MFG | 38.95 | *** | 43.66 | *** |

Notes: All *p* values were corrected by FDR. *$*P < 0.05$; $**P < 0.01$; $***P < 0.001$*; n.s., no significance. CMA, centromedian amygdala; BLA, basolateral amygdala; mCC, middle portions of cingulate cortex;

PCC, posterior portions of cingulate cortex; VC, visual cortex; ITC, inferior temporal cortex; MTG, middle temporal gyrus; STG, superior temporal gyrus; PMC, premotor cortex; SM, sensorimotor cortex; ACC, anterior cingulate cortex; IFG, inferior frontal gyrus; MPFC, medial prefrontal cortex; MFG, middle frontal gyrus.

**Table. S3. Brain regions showing significant difference in functional connectivity difference (BLA versus CMA) between integration versus segregation (Cohort 1)**

| Brain regions | | State 1 | | State 2 | |
|---|---|---|---|---|---|
| | | BLA versus CMA | | | |
| | | T values | *p* values | T values | *p* values |
| Limbic and paralimbic structures | Insula | -40.59 | *** | -41.60 | *** |
| | Hippocampus | 102.19 | *** | 107.48 | *** |
| | mCC | 21.78 | *** | 21.69 | *** |
| | PCC | 31.72 | *** | 35.22 | *** |
| Uni- and polymodal association cortex | VC | 19.41 | *** | 26.09 | *** |
| | Parahippocampal gyrus | 69.22 | *** | 72.19 | *** |
| | ITC | 54.51 | *** | 57.81 | *** |
| | MTG | 52.59 | *** | 57.26 | *** |
| | STG | 58.68 | *** | 68.16 | *** |
| | PMC | 35.70 | *** | 39.10 | *** |
| | SM | 41.34 | *** | 43.67 | *** |
| Subcortical structures | Stratum | -69.15 | *** | -70.18 | *** |
| | Thalamus | -40.74 | *** | -34.90 | *** |
| | Midbrain | 36.55 | *** | 34.03 | ** |
| Cerebellum | Vermis/declive | 39.40 | *** | 39.02 | *** |
| | Crus | -41.31 | *** | -38.48 | *** |
| Prefrontal cortex | ACC | 5.29 | *** | 12.48 | *** |
| | IFG | 12.81 | *** | 14.56 | *** |
| | MPFC | 32.68 | *** | 42.92 | *** |
| | MFG | 1.33 | n.s. | 3.93 | *** |

Notes: All notes and abbreviations are the same as that in Table S2.

**Table. S4. Coefficients of brain regions for predicting SCLs during integration state (Cohort 2)**

| Regions of interest | | BLA-target | CMA-target |
|---|---|---|---|
| | | Weight | Weight |
| Limbic and paralimbic structures | Insula | 0.305 | -0.518 |
| | Hippocampus | -0.242 | 0.000 |
| | mCC | -0.649 | 0.406 |
| | PCC | 0.387 | -0.333 |
| | VC | -0.045 | 0.106 |
| | Parahippocampal gyus | 0.243 | 0.000 |
| Uni- and polymodal association cortex | ITC | 0.086 | 0.668 |
| | MTG | 0.000 | 0.000 |
| | STG | -0.314 | -0.475 |
| | PMC | 0.189 | 0.566 |
| | SM | -0.507 | 0.254 |
| Subcortical structures | Stratum | 0.480 | -0.385 |
| | Thalamus | -0.375 | 0.542 |
| | Midbrain | -0.149 | 0.087 |
| Cerebellum | Vermis/declive | 0.029 | 0.084 |
| | Crus | -0.081 | -0.124 |
| Prefrontal cortex | ACC | -0.041 | -0.528 |
| | IFG | 0.264 | 0.557 |
| | MPFC | 0.220 | -0.252 |
| | MFG | 0.000 | -0.288 |

Notes: Weight is the contribution of the functional connectivity between BLA and CMA. Other notes and abbreviations are the same as that in Table S2.

**Table. S5. Coefficients of brain regions for predicting SCLs during segregation state (Cohort 2)**

| Regions of interest | | BLA-target | CMA-target |
|---|---|---|---|
| | | Weight | Weight |
| Limbic and paralimbic structures | Insula | 0.000 | -0.143 |
| | Hippocampus | -0.361 | 0.099 |
| | mCC | 0.066 | -0.074 |
| | PCC | -0.175 | -0.249 |
| Uni- and polymodal association cortex | VC | 0.000 | -0.160 |
| | Parahippocampal gyrus | 0.202 | 0.258 |
| | ITC | 0.000 | -0.271 |
| | MTG | 0.000 | -0.009 |
| | STG | -0.018 | 0.000 |
| | PMC | -0.114 | 0.019 |
| | SM | -0.234 | 0.058 |
| Subcortical structures | Striatum | 0.150 | -0.027 |
| | Thalamus | -0.087 | 0.085 |
| | Midbrain | 0.000 | 0.289 |
| Cerebellum | Vermis/declive | -0.120 | 0.275 |
| | Crus | 0.000 | -0.026 |
| Prefrontal cortex | ACC | -0.534 | -0.137 |
| | IFG | -0.141 | 0.254 |
| | MPFC | 0.595 | 0.058 |
| | MFG | 0.334 | -0.103 |

Notes: Weight is the contribution of the functional connectivity between BLA and CMA. Other notes and abbreviations are the same as that in Table S2.
